# Simple, sensitive, and cost-effective detection of wAlbB *Wolbachia* in *Aedes* mosquitoes, using loop mediated isothermal amplification combined with the electrochemical biosensing method

**DOI:** 10.1101/2021.06.30.450550

**Authors:** Parinda Thayanukul, Benchaporn Lertanantawong, Worachart Sirawaraporn, Surat Charasmongkolcharoen, Thanyarat Chaibun, Rattanalak Jittungdee, Pattamaporn Kittayapong

## Abstract

**Background:** *Wolbachia* is an endosymbiont bacterium generally found in about 40% of insects, including mosquitoes, but it is absent in *Aedes aegypti* which is an important vector of several arboviral diseases. The evidence that *Wolbachia* trans-infected *Ae. aegypti* mosquitoes lost their vectorial competence and became less capable of transmitting arboviruses to human hosts highlights the potential of using *Wolbachia-* based approaches for prevention and control of arboviral diseases. Recently, release of *Wolbachia* trans-infected *Ae. aegypti* has been deployed widely in many countries for the control of mosquito-borne viral diseases. Field surveillance and monitoring of *Wolbachia* presence in released mosquitoes is important for the success of these control programs. So far, a number of studies have reported the development of loop mediated isothermal amplification (LAMP) assays to detect *Wolbachia* in mosquitoes, but the methods still have some specificity issues.

**Methodology/Principal Findings:** We describe here the development of a LAMP combined with the DNA strand displacement-based electrochemical sensor (BIOSENSOR) method to detect wAlbB *Wolbachia* in trans-infected *Ae. aegypti*. Our developed LAMP primers were more specific to wAlbB detection than those of the previous published ones if the assays were conducted with low-cost and non-specific detecting dyes. The detection capacity of our LAMP technique was 3.8 nM and the detection limit reduced to 2.16 fM when combined with the BIOSENSOR. Our study demonstrates that the BIOSENSOR can also be applied as a stand-alone method for detecting *Wolbachia*; and it showed high sensitivity when used with the crude DNA extracts of macerated mosquito samples without DNA purification.

**Conclusions/Significance:** Our results suggest that both LAMP and BIOSENSOR, either used in combination or stand-alone, are robust and sensitive. The methods have good potential for routine detection of *Wolbachia* in mosquitoes during field surveillance and monitoring of *Wolbachia*-based release programs, especially in countries with limited resources.

**Author Summary:** Mosquito-borne diseases such as dengue, chikungunya, Zika, and yellow fever are transmitted to humans mainly by the bites of *Aedes aegypti* mosquitoes. Controlling these diseases relies mostly on the use of insecticides, in which the efficiency has been reduced through development of insecticide resistance in mosquitoes. *Wolbachia* is the endosymbiotic bacteria that are naturally found in 40% of insects, including mosquitoes. The bacteria could protect their hosts from viral infections and could also cause sterility in host populations, therefore, providing an opportunity to use them for disease control. Application of a *Wolbachia*-based strategy needs simple, rapid and sensitive methods for detecting the bacteria in released mosquitoes. In this paper, we develop the combined methods of LAMP and BIOSENSORS for detecting wAlbB *Wolbachia* in mosquitoes. Our positive LAMP reaction can be visualized by color change from violet to blue at a sensitivity of ≥ 60 pg of genomic DNA. When used in combination with the BIOSENSOR method, the sensitivity increased a million fold without losing specificity. Our study indicates that both developed methods, either used in combination or stand-alone, are efficient and cost-effective, hence, it could be applied for routine surveys of *Wolbachia* in mosquito control programs that use *Wolbachia*-based approaches.

## Introduction

Dengue, chikungunya, Zika, and yellow fever diseases, transmitted by the *Aedes aegypti* vector, continue to be a major health problem and affect human populations worldwide. Prevention of the transmission of these diseases, when vaccines have not yet been fully effective, depends primarily on two approaches, i.e., mosquito control and interruption of human-vector contact [1]. Historically, insecticides have been the primary means of mosquito control; however, the overuse and misuse of insecticides has resulted in several negative consequences. The use of many of these insecticides are problematic today; these problems include deleterious impacts on the environment and the emergence of insecticide-resistant mosquitoes [2]. Alternative vector control strategies are important and need to be considered to effectively control the spread of vector-borne diseases.

*Wolbachia* is an endosymbiont found intracellularly in about 40% of insect species [3]. The bacteria can manipulate host reproduction and inhibit virus intracellular replication; hence it is potentially an effective alternative to traditional chemical pesticides. In mosquitoes, *Wolbachia* can induce cytoplasmic incompatibility (CI), a phenotype which results in the production of unviable offspring when uninfected females mate with *Wolbachia*-infected male mosquitoes. On the other hand, if *Wolbachia*-infected females mate with either infected or uninfected male mosquitoes, viable progenies harboring maternally transmitted *Wolbachia* will be produced. The effect of CI has received much attention, as it offers the potential application of *Wolbachia* in vector control. There have been a number of reports describing the stable establishment of *Wolbachia* in mosquitoes [5–7]. The use of *Wolbachia*-based approaches to reduce transmission of dengue, Zika, and other *Aedes*-borne disease viruses is currently being deployed and implemented widely in international programs in many countries [8, 9].

Although large-scale release of *Wolbachia* trans-infected *Ae. aegypti* populations into the wild has been occuring in many countries, there remains critical issues to be addressed with respect to this strategy in order to maintain the quality of the released mosquitoes. Surveillance of mosquito infection status is critical for the planning and deployment of proper mosquito control initiatives. Thus far, PCR has been the gold standard method used for detecting *Wolbachia* in mosquitoes [10, 11]. However, the method is laboratory based, requires trained personnel, and uses expensive instruments. Subsequently, loop-mediated isothermal amplification (LAMP), a highly sensitive and specific amplification of target DNA, was developed and is used for detecting *Wolbachia* in *Ae. aegypti*. To detect a diverse range of *Wolbachia* strains, LAMP primer sets were developed based on the 16S rRNA gene [12, 13]. To evaluate the efficacy of the *Wolbachia* trans-infected mosquito interventions, LAMP primers specific to wAlbB and wMel strains were developed based on *Wolbachia* surface protein gene (wsp) [14, 15]. High fidelity detection using LAMP combined with oligonucleotide strand displacement (OSD) probes, and enhancement of the LAMP reaction speed using two loops, have been developed [14, 16]. Recently, in Northern Australia, trials releasing wAlbB-infected *Ae. aegypti* were implemented [17]. *Wolbachia* wAlbB infected *Ae. aegypti* is suitable to apply for mosquito control in hot climate regions, because *Wolbachia* density remains high at 26–40°C, which is in contrast to wMel and wMelPop infected mosquitoes [18]. Therefore, the LAMP detection for the wAlbB-infected mosquitoes should be further developed and validated, so as to establish a robust, sensitive, specific detection of *Wolbachia* in field released *Ae. aegypti* mosquitoes.

The LAMP products can be analyzed either by agarose gel electrophoresis or visual inspection of color or turbidity changes [19]. Therefore, the disadvantage of the method is mis-diagnosis caused by a false positive or false negative. An alternative method to overcome the problem is the use of electrochemical-DNA based biosensor, which employs gold-nanoparticles (AuNPs) to electrochemically label nucleic acid [20–23]. In this paper, we describe the development of a combined LAMP and electrochemical-DNA based biosensor with the strand displacement reaction methods in order to detect wAlbB *Wolbachia* trans-infected *Ae. aegypti* mosquitoes.

## Methods

### Ethical issues

The use of mosquito colony materials in this study was approved by the Faculty of Science, Mahidol University Animal Care and Use Committee (SCNU-ACUC) (Protocol No. MUSC64-005-554).

### Mosquito materials and genomic DNA extraction

Laboratory rearing and field collected mosquitoes including *Aedes aegypti* (Aae-JJ), *Aedes albopictus*(Aal-CH), wAlbB trans-infected Thai *Aedes aegypti* (wAlbB-TH), and *Culex quinquifasciatus* (Cq-BK) were used in this study. The wAlbB trans-infected Thai *Aedes aegypti* was generated using the direct microinjection technique as previosly described [5, 24]. The mosquito genomic DNA was extracted using the crude boiling method [25]. Briefly, mosquito samples were ground in 100 μl of Sodium Chloride-Tris-EDTA buffer (STE; 10 mM Tris-HCl pH 8.0, 1 mM EDTA and 100 mM NaCl), heated for 10 min at 95°C, and centrifuged. Supernatant was transferred to a new tube and used as a template sample in subsequent LAMP, PCR, and biosensor reactions.

### LAMP primers and biosensor probe design

The sequence of the *wsp* gene of wAlbB trans-infected Thai *Ae. aegypti* (MZ325222) was applied for designing the LAMP primers. The sequence was identical to AF020059 wAlbB from *Aedes albopictus* (Houston strain) and MN307069 *Wolbachia* of *Aedes aegypti* isolate wAegB from NCBI GenBank. This sequence was submitted to Primer Explorer v5 software (primerexplorer.jp/lampv5e/index.html, Eiken Chemical Co., Japan) to generate the potential primers used in the wAlbB LAMP detection. Several potential LAMP primer sets were generated. The highly recommended regions were compared to various *wsp* sequences in the NCBI GenBank database. The DNA alignment was performed using MEGA 7.0.26 software [26]. The consensus regions among most wAlbB were used to construct LAMP primers and biosensor probes (Table 1). All primers and probes were synthesized by Bio Basic Canada, Inc. Canada and Integrated DNA Technologies, USA, respectively.

**Table 1.**
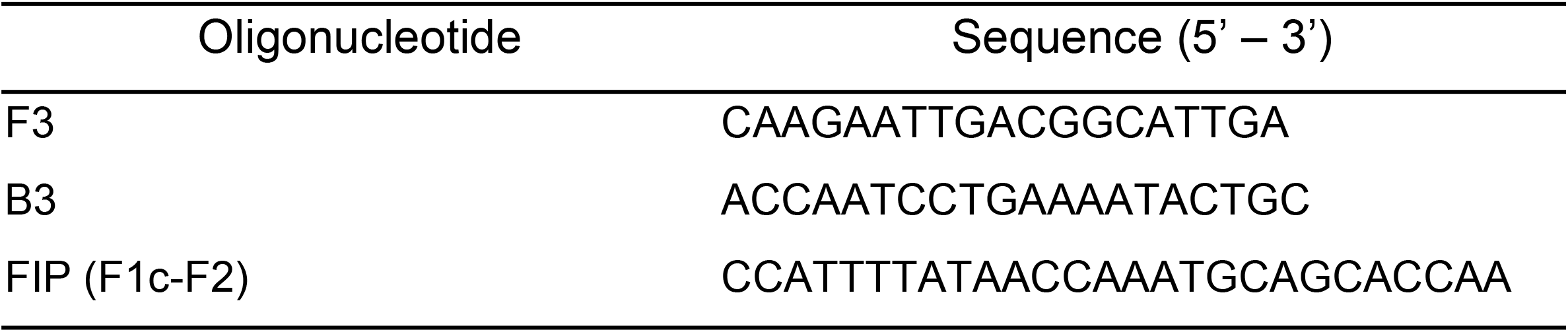
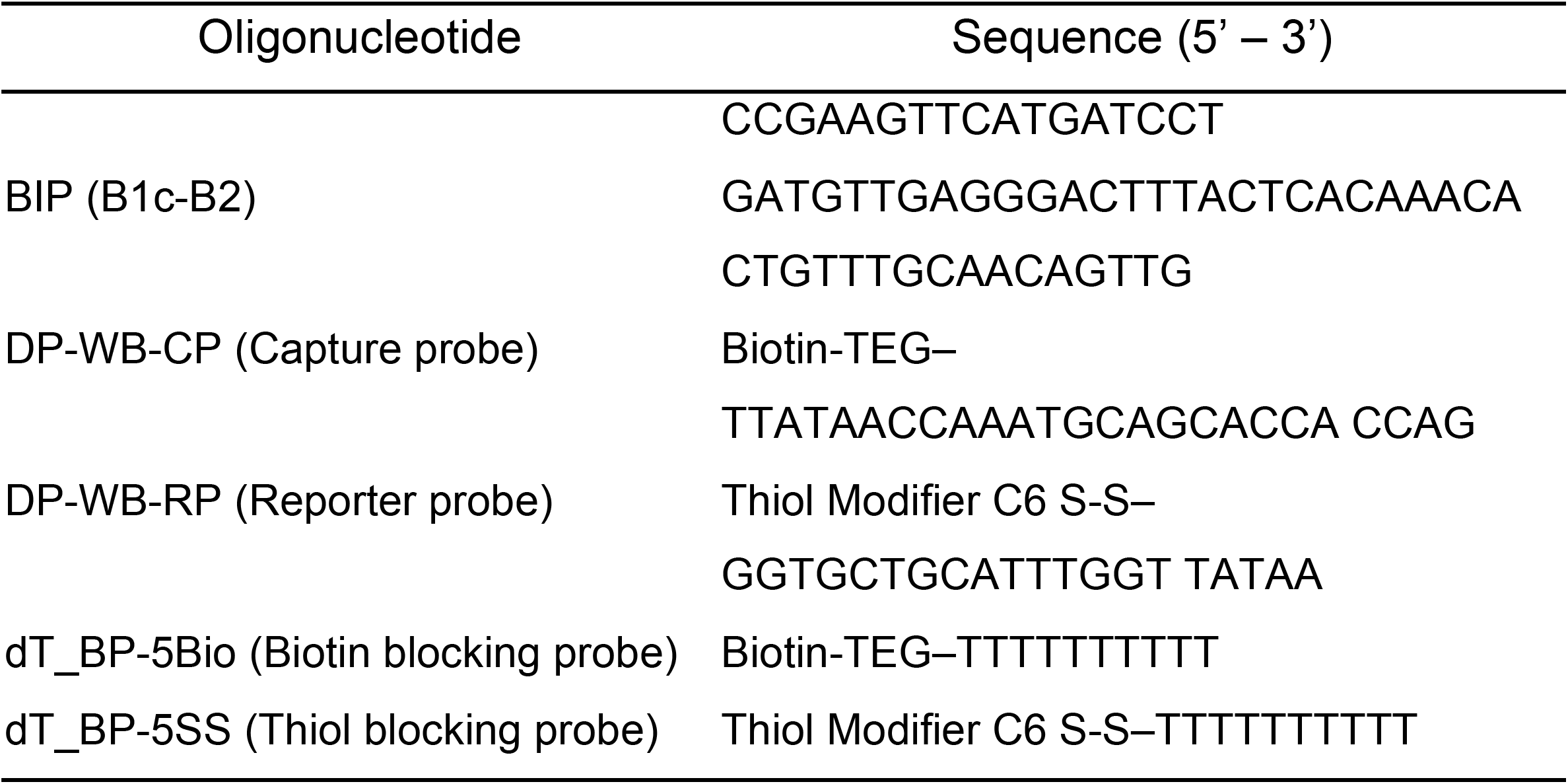
Oligonucleotide sequences of LAMP primers and target induced-DNA strand displacement probes used in this study.

### LAMP reaction

LAMP assay was performed in a total volume of 25 μl with *Bst* 2.0 WarmStart™ DNA Polymerase (New England Biolabs). The reagents, modified from [14], consisted of 1× Isothermal buffer (20 mM Tris-HCl, 10 mM (NH_4_)_2_SO_4_, 50 mM KCl, 2 mM MgSO_4_, 0.1% Tween 20, pH 8.8 at 25°C), 0.4 mM of each dNTPs (10 mM each, Invitrogen, USA), 0.8 M Betaine solution (5 M, Sigma, USA), 2 mM MgSO_4_, 1.6 μM of each internal primer (FIP/BIP), 0.4 μM of each external primer (F3/B3), 6.4 units of *Bst* 2.0 DNA polymerase, 120 μM of hydroxy naphthol blue (HNB), metal (pM) indicator, Merck (Germany), and 4 μl DNA or approximately 20 ng of DNA. HNB is a visualizing indicator of magnesium ion reduction due to the magnesium pyrophosphate formation by LAMP [20]. The mixture was incubated at 65°C for 90 min, followed by 80°C for 10 min. The concentrations of DNA (20–100 ng), *Bst* 2.0 (1.6–8.0 U), and the reaction time (60–90 min) were varied depending upon experimental purposes as indicated.

### PCR reaction and gel electrophoresis

*Wolbachia* detection was performed using primers wsp 81F and 691R for general *Wolbachia* detection (~600 bp); primers 183F and 691R for wAlbB detection (~500 bp); and primers 328F and 691R for wAlbA detection (~380 bp), according to the method previously described [27]. The reactions were performed using final volumes of 25 μl, including 1.25 U *Taq* recombinant DNA polymerase (Invitrogen, USA), 1x PCR Buffer (-Mg), 3.75 mM MgCl_2_, 0.25 mM each dNTP (Invitrogen, USA), 0.5 μM each primer, and 1.0 μl template DNA. The amplifications were performed using a thermal cycler (T100™ Thermo Cycler, Biorad, USA) with the following parameters: one step of 3 min at 94°C, 35 cycles of 45 s at 94°C, 30 sec at annealing temp (wsp 58°C; Den 55°C, 16S 53°C), 45 sec at 72°C, and one step of 10 min at 72°C. All PCR products were detected by electrophoresis on a 2.0% (w/v) Agarose A gel (Biobasic, Canada)) containing 0.2 μg/ml Ethidium Bromide (Sigma, USA) in 1xTBE buffer (pH 8.0) at 100 V for 40 min and visualized under UV light. Five μl of the PCR and LAMP product was mixed with 2 μl of loading dye. Nucleic acid concentration was measured using NanoDrop™ One Microvolume UV-Vis Spectrophotometer (Thermo Fisher Scientific, USA)

### Functionalization of AuNPs conjugate with reporter probe

Preparation of AuNPs–reporter probe (AuRP) conjugate was performed using the salt aging method [24]. Briefly, 10 μl of 100 μM reporter probe DNA and 30 μl of 100 μM blocking probe sequences (PolyT_10_) thiolated DNA were activated by using 10 mM Tris (2-carboxyethtl) phosphine (TCEP, Sigma-Aldric, USA) freshly prepared. Then, the thiol-activated DNA was added into 1 ml of 40 nm AuNPs solution (DCN Diagnostics, USA) and incubated overnight at room temperature. After incubation, the solution of 10 μl of 500 mM Tris-acetate pH 8.2 and 1 M NaCl 100 μl was added into the mixture and stored overnight before the next step. The excess probes were isolated by centrifugation at 14,000 rpm 30 min, followed by washing 3 times with 25 mM Tris-acetate pH 7.4, resuspension with hybridization buffer, and then stored at 4°C until use.

### Immobilizations capture probe on magnetic bead particle

The immobilization of the biotinylated capture probe (CP) on the magnetic bead (MB), (Dynabeads T1, Thermo Fisher Scientific, USA) was performed according to the manufacturer’s instruction. A 100 μl (10 μg/μl) of MB was washed 3 times with 200 μl of 20 mM PBS pH 7.4, mixed with 4 μl of 100 μM capture probe, 12 μl of 100 μM Biotin probe and 184 μl of 20 mM PBS pH 7.4, and then incubated for 40 min at room temperature. The MB-bound probe was washed 3 times with 20 mM PBS pH 7.4, resuspended with 100 μl of 20 mM PBS pH 7.4, and then stored at 4°C until use. This conjugation was subsequently called magnetic bead conjugated capture probe DNA (MB-CP).

### DNA hybridization and DNA strand displacement reaction

The prehybridization step of MB-CP and AuRP was prepared as follows: 2 μl of MB-CP and 10 μl of AuRP were added into 18 μl of 20 mM PBS/0.1% SDS pH 7.4, and then incubated for 20 min at 45°C in a water bath. The prehybridized MB-CP and AuRP was then washed 3 times with 20 mM PBS pH 7.4 using magnet collection. The pellet was used for a DNA strand displacement experiment. For DNA strand displacement, 30 μl of target DNA was added to resuspend the pellet and then incubate at 60°C for 30 min. A magnet was used to separate the target DNA bound to MB-CP from the unbound (displaced). AuNP-RP and 5 ml of the supernatant was used for signal detection.

### Electrochemical detection of AuRP from DNA strand displacement reaction

Displaced AuRP was detected by using electrochemical measurement by the differential pulse anodic stripping voltammetry (DPASV) technique on Palmsens 4 computer-controlled potentiostat with PSTrace version 5.7 software (Palmsens, The Netherlands). Two electrode systems screened printed carbon electrodes or SPCE (Quesence, Thailand) — which consisted of two carbon tracks as working electrode, reference electrode, and counter electrode in DPASV — were used. 5 μl of the desired sample was loaded onto a working electrode surface, followed by 50 μl of 1 M hydrobromic acid (HBr)/0.1M bromine solution (Br_2_). For the pre-treatment step, the condition for deposition potential was −0.75 V and the deposition time was 100 sec. The step potential was set at 0.005 V, with the interval time set at 0.1 sec. The modulation amplitude was 0.1 V and the modulation time 0.05 sec.

### Detection of nucleic acid derived from wAlbB

The products of the wAlbB LAMP reaction and PCR of *wsp* genes, wAlbA *wsp* gene, and wAlbB *wsp* gene were applied to the biosensor detection. In addition, the macerated mosquito samples from the laboratory colony and field collection were used in this study. The concentration of the DNA was determined by measuring the absorbance at 260 nm, using the NanoDrop™ One Microvolume UV-Vis Spectrophotometer (USA) in the DNA strand displacement platform, followed by differential pulse anodic stripping voltammetry (DPASV) detection.

## Results

### LAMP primer and probe design

The *wsp* genes of *Wolbachia* trans-infected Thai *Ae. aegypti* were sequenced. This sequence was compared to 17 *wsp* genes of *Wolbachia* wAlbB in mosquitoes from the NCBI database. The consensus region around 230 bp was submitted to PrimerExplorer software. The recommended LAMP primer sets were compared to 686 sequences of *wsp* genes from 66 *Wolbachia* strains [27–29] and all *wsp* genes of *Ae. aegypti* and *Ae. albopictus* in the database. The set of sequences that could bind to all wAlbB sequences and were different from most other strains were selected (Table 1).

By comparing the *wsp* gene sequences and considering that LAMP may amplify even with few primer bindings, it was seen that our primers could have non-specific bindings to some *Wolbachia* strains in mosquitoes, including wPip in *Ae. aegypti* (MK860184-5), *Ae. albopictus* (MF805773, MF805775) and *Cx. pipiens quinquefasciatus* (AF301012); wAnsA in *Anopheles* sp. (MH605284); *Wolbachia* strains in some *Cx. quinquefasciatus* (KJ140125), *Cx. tritaeniorhynchus* (KY457713), *Cx. pipiens* (KJ500030); wFus in *Cx. fuscocephala* (AF317481); wMad in *Aedeomyia madagascarica* (MK033272); wKes in *Armigeres kesseli* (AF317489); *Wolbachia* in some *Ar. subalbatu*s (KY457714, KY457720) and *Ar. obturbans* (KJ140130, KJ140132), and wPseu in *Ae. pseudalbopictus* (AF317487). The nonspecific bindings of the wAlbB specific LAMP primers in the previous study [14] were broader, as they could bind more *Wolbachia* strains in mosquitoes, including wDec in *Cx. decens* (MK033274); wUra2 in *Uranotaenia* sp. (MK033278); wUnif in *Mansonia uniformis* (AF317493); some *Wolbachia* in *Ma. uniformis* (MH777433, KY523674); wNoto in *Ae. notoscriptus* (KT962260); wFlu in *Ochlerotatus fluviatilis* (KF898395); wInd in *Ma. Indiana* (AF317492); wSit in *Cx. sitiens* (AF317491); wPerp in *Ae. perplexus* (AF317486); wGel in *Cx. gelidus* (AF317482); and wCra in *Coquillettidia crassipes* (AF317478) but they could not bind with wAnsA in *Anopheles* sp. (MH605284).

Since the previous work [14] used oligonucleotide strand displacement (OSD) probes and quencher technology for LAMP detection, the assay reported only the probe priming region. The ‘WSP-OSD’ probe in the previous work [14] could detect wPip (MK860184-5, MF805775), *Wolbachia* strains in some *Culex* spp. (AF301012, KJ140125, KY457719, AF317487); *Armigeres* spp. (KY457714, KY457720, KJ140132, KJ140130, AF317489), and *Aedeomyia madagascarica* (MK033272), whereas the ‘wAlbB vs wPip OSD probe’ [14] or ‘WSP.BLP loop’ reported in another study [16] would bind only wAlbB, *Wolbachia* strains in some *Armigeres* spp. (KY457720, KJ140132, KJ140130, AF317489); and wPseu in *Aedes pseudalbopictus* (AF317487). If the LAMP assays were conducted with non-specific detecting methods like HNB dye, Sybr green I, or Cresol Red, our LAMP primer set would be more specific to wAlbB detection.

In addition, the new primers B3, FIP, and BIP in our study had higher GC rate (40-42%), closer to the recommended range for good binding primers of 50-60% [30] than those reported previously (35-40%). Our LAMP primers had a melting temperature in the range of 55.2–61.3°C, where the delta G values of 3’ and 5’ ends were −6.24 to −4.07 kcal/mol and −5.69 to −4.02 kcal/mol, respectively, and the dimer (minimum) delta G was −2.16 kcal/mol. For the capture probe design, we selected the consensus region overlapping with the F1c binding area, so as to increase the attachment of the probe to the structures of complex LAMP products (Figure 1).

**Figure 1.**
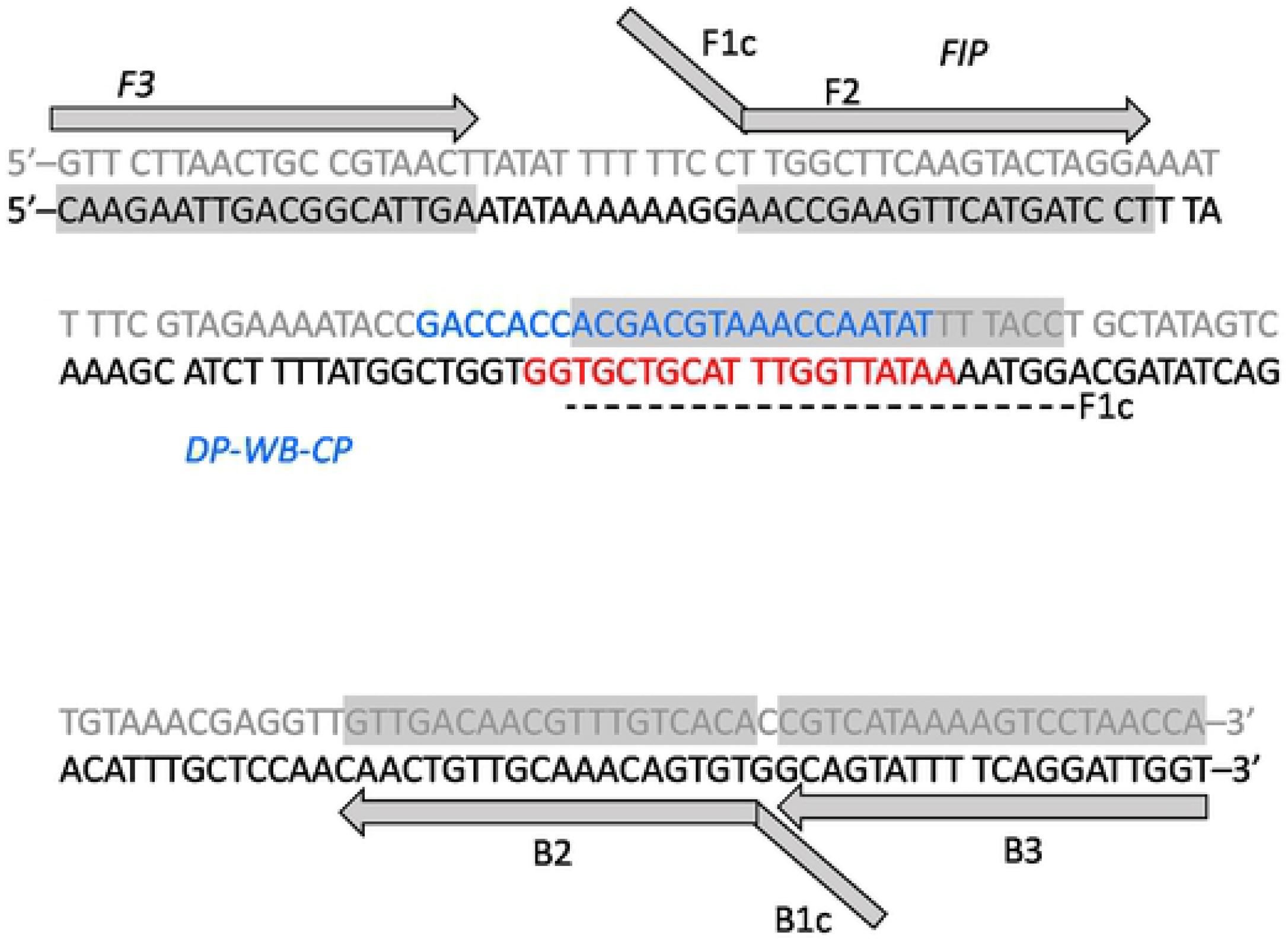
Schematic diagram demonstrates the LAMP primers and probe binding locations on the target sequences of the wAlbB *wsp* gene. Grey boxes indicate the primer sequences. Grey letters are the complementary sequence of the wAlbB sequence in the 5’-3’ direction. Blue and red fonts indicate the capture and reporter probes, respectively.

### LAMP assay

The developed LAMP primer set was used to examine the presence of *Wolbachia* in mosquito samples, i.e., *Ae. albopictus* naturally superinfected with wAlbA and wAlbB, *Cx. quinquefasciatus* naturally infected with wPip [29], wild-type *Ae. aegypti* mosquitoes which do not harbor *Wolbachia* [31], and wAlbB trans-infected Thai *Ae. aegypti* (Figure 2). The LAMP assay clearly showed positive *Wolbachia* detection for the *Wobachia* trans-infected Thai *Ae. aegypti*, *Ae. albopictus*, and *Cx. quinquefasciatus*, as indicated by the blue color of HNB in the reactions due to the loss of Mg^2+^ ions to the magnesium pyrophosphate precipitation and the presence of a ladder-like band pattern upon gel electrophoresis, while wild-type *Ae. aegypti* and the control reaction (no-template control (NTC)), which were negative, gave a purple color without ladder-like bands that was observed upon gel electrophoresis. These results were in agreement with the gold standard PCR method, suggesting the potential of these newly designed LAMP primers to detect *Wolbachia* infection in mosquitoes. It is noteworthy that, as predicted from *in silico* analysis, all primers except for B3 could bind to the wsp sequence of the laboratory *Cx. quinquefasciatus* (MZ325223). The previous LAMP-OSD assay also detected wPip in *Cx. quinquefasciatus* in their study. To verify the efficiency on differentiating wAlbB strains, we repeated the test with 45 *Ae. albopictus*, 20 wAlbB trans-infected Thai *Ae. aegypti*, and 20 wild-type *Ae. aegypti*. All tests gave the expected results correctly. It is noteworthy that LAMP could amplify the DNA binding target even though not all primers bound to the region, as shown in the case of *Cx. quinquefasciatus*.

**Figure 2.**
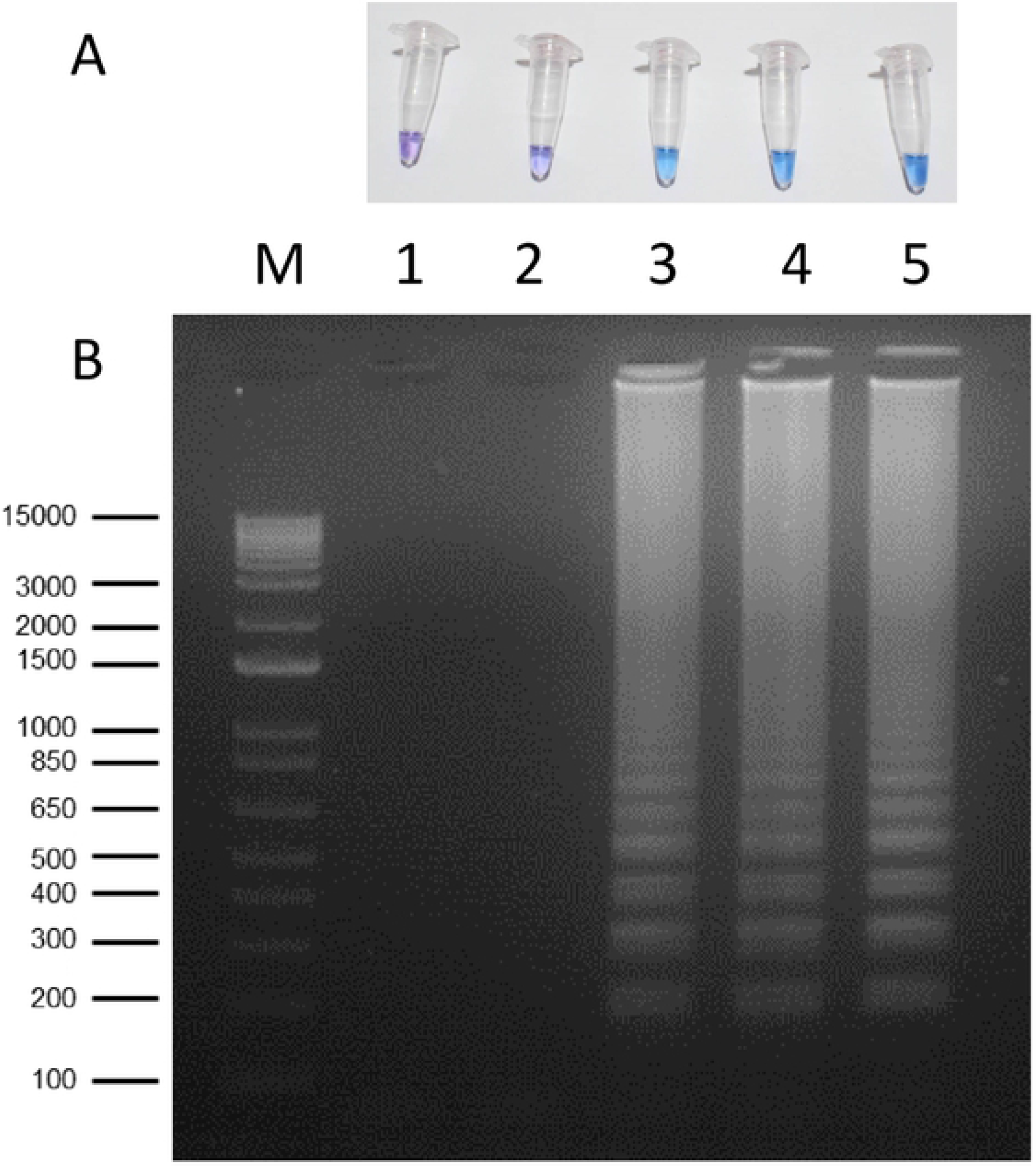
Detection of *Wolbachia* wAlbB gene in different mosquito species using LAMP assay with HNB indicator (A) and ethidium bromide stained gel (B): (1) no template control, (2) wild-type *Aedes aegypti*, (3) *Wobachia* trans-infected Thai *Aedes aegypti*, (4) *Aedes albopictus*, and (5) *Culex quinquefasciatus*. M is 1 kb plus DNA Ladder from Invitrogen™. The condition for LAMP reaction was 6.4 units of *Bst* 2.0 DNA polymerase, 65°C for 60 min and 80°C for 10 min.

To optimize LAMP detection, the concentration of the DNA template was examined. *Ae. albopictus* and wAlbB trans-infected Thai *Ae. aegypti* were used as positive controls. Wild type *Ae. aegypti* and NTC were used as negative controls. The amounts of DNA template varied between 20 and 100 ng (supplementary Figure S1). The ladder-like bands were observed for *Ae. albopictus* and wAlbB trans-infected Thai *Ae. aegypti* samples for all template amounts, which were in contrast to the results of wild-type *Ae. aegypti* and NTC. Wild-type *Ae. aegypti* showed a darker blue-purple color closer to the positive control at a higher amount of DNA. However, at the DNA amount of 100 ng, the color from the wild-type *Ae. aegypti* reaction could not be differentiated from that of the positive reaction. Therefore, the amount of DNA should be controlled in the range of 20–80 ng, with the most recommended DNA amount being 20 ng.

The LAMP assay was tested with different *Bst* polymerase concentrations from 1.6–8.0 Units (Supplementary Table S1). The color development between the positive and negative control could be more distinguished at higher concentrations of *Bst* (3.2–8.0 Units). At 1.6 Units *Bst*, a false negative result was obtained for the *Wolbachia* trans-infected Thai *Ae. aegypti*. Hence, *Bst* at concentrations of 3.2–6.4 Units were recommended for cost saving and visual observation. However, in the present study, we used 6.4 Units of *Bst*, as was done in many previous studies [12, 14, 32].

We also tried to develop the LAMP reaction at 60 and 90 min for 20 and 40 ng template DNA (Figure S2). The HNB and gel results were positive for *Ae. albopictus* (P) and negative for NTC at both 60 and 90 min for 20–40 ng DNA. However, the blue lavender color in the 60-min reaction of *Ae. albopictus* at 40 ng was ambiguous for visual observation. Therefore, 90 min was recommended. This is consistent with previous studies which also suggested 90 min for LAMP amplification [12, 14]. Besides, the concentration of HNB used in this study was only 0.12 mM, which is 10-time less than that reported in the previous work [12]. However, the range of HNB dye concentration could be varied without affecting the LAMP reaction.

The LAMP reaction was performed with diluted DNA from the *Ae. albopictus* sample (156.6 ng/μl) in a total reaction volume of 25 μl with 4 μl DNA template. A 10-fold dilution of the DNA sample was prepared (Figure 3). Both LAMP and PCR could amplify the positive samples up to 10^−4^ dilution, which was equivalent to 62.6 pg DNA in 25 μl, or a DNA concentration of 3.8 nM. However, both LAMP and PCR techniques failed to detect DNA at a concentration below 0.38 nM. At a DNA concentration 3.8 nM, PCR yielded a very faint band, whereas color visualization and ladder-bands of LAMP reaction could be distinguishable from the negative control reaction. Therefore, LAMP had better sensitivity than PCR. We concluded that the limit of detection (LOD) of LAMP in this study was 3.8 nM.

**Figure 3.**
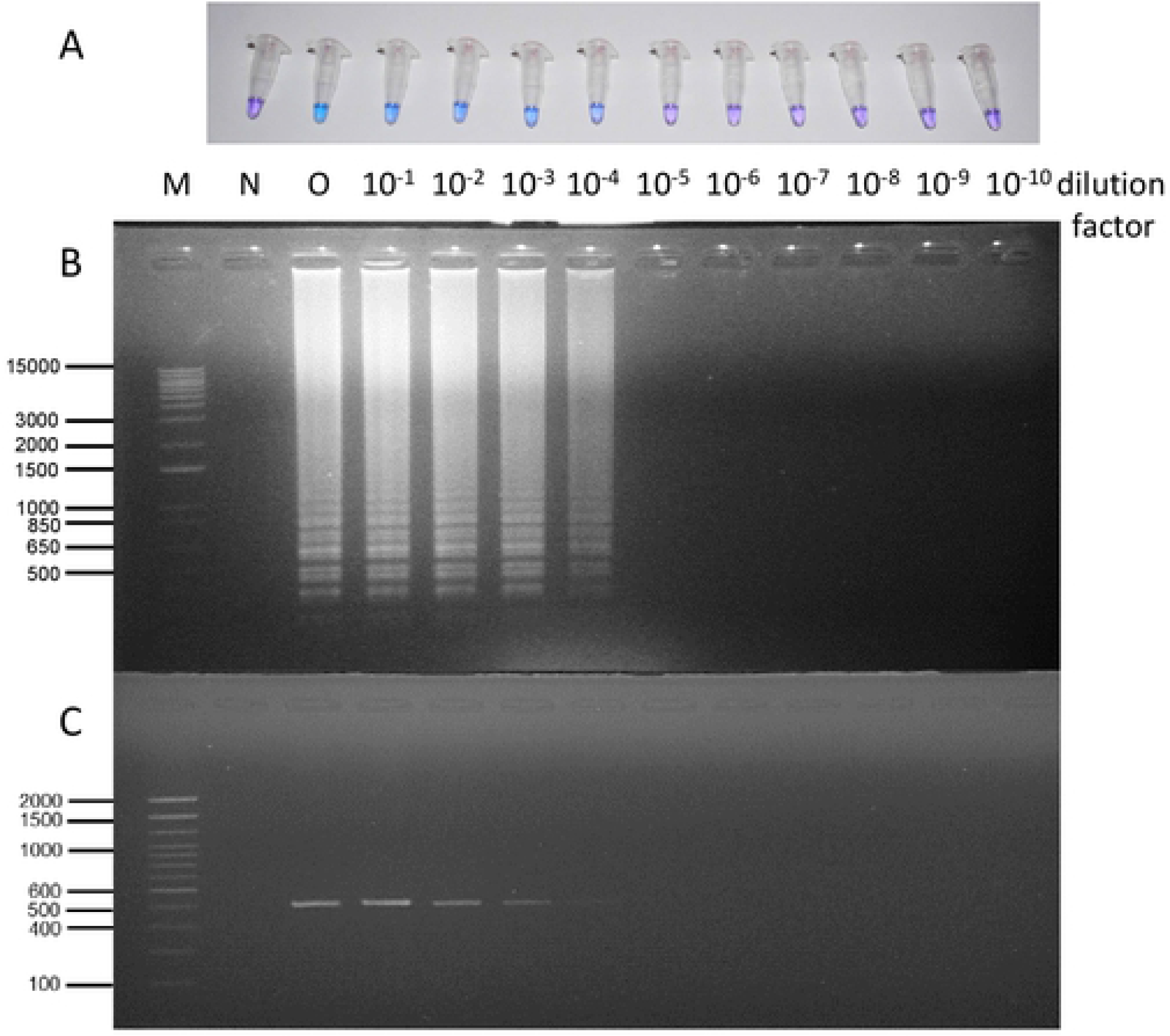
Detection of *Wolbachia* wAlbB gene using LAMP assay with HNB indicator (A) and ethidium bromide stained gel (B) of a 10-fold dilution of an individual *Aedes albopictus* sample (156.6 ng/μl, 260/280 = 1.8, 260/230 = 0.89) including 10^1^– 10^−10^ times. Polymerase chain reaction with wsp primers (691R and 183F) of the same dilution (C). (6.4 units of *Bst* 2.0 DNA polymerase, 65°C for 60 min and 80°C for 10 min) (O) is non-diluted original sample, (N) is No template control, (M upper) is Invitrogen™ 1 kb plus DNA Ladder, and (M lower) is Invitrogen™ 100 bp DNA Ladder.

### DNA sensors assay

Biosensor was applied to increase the sensitivity of LAMP detection and to reduce ambiguity in LAMP visualization. Figure 4 shows the sensitivity of the strand displacement method with a synthetic wAlbB linear target using electrochemical detection. The results showed the LOD of 2.16 fM for the target DNA (5 Signal/Noise). The linear range was 1 fM to 1 μM (R^2 =^ 0.93). The electrochemical with the target strand displacement platform has a much higher sensitivity than the LAMP and PCR techniques in a magnitude of 10^6^. Therefore, the sensitivity of *Wolbachia* DNA detection could be enhanced dramatically by using the electrochemical DNA sensors technology.

**Figure 4.**
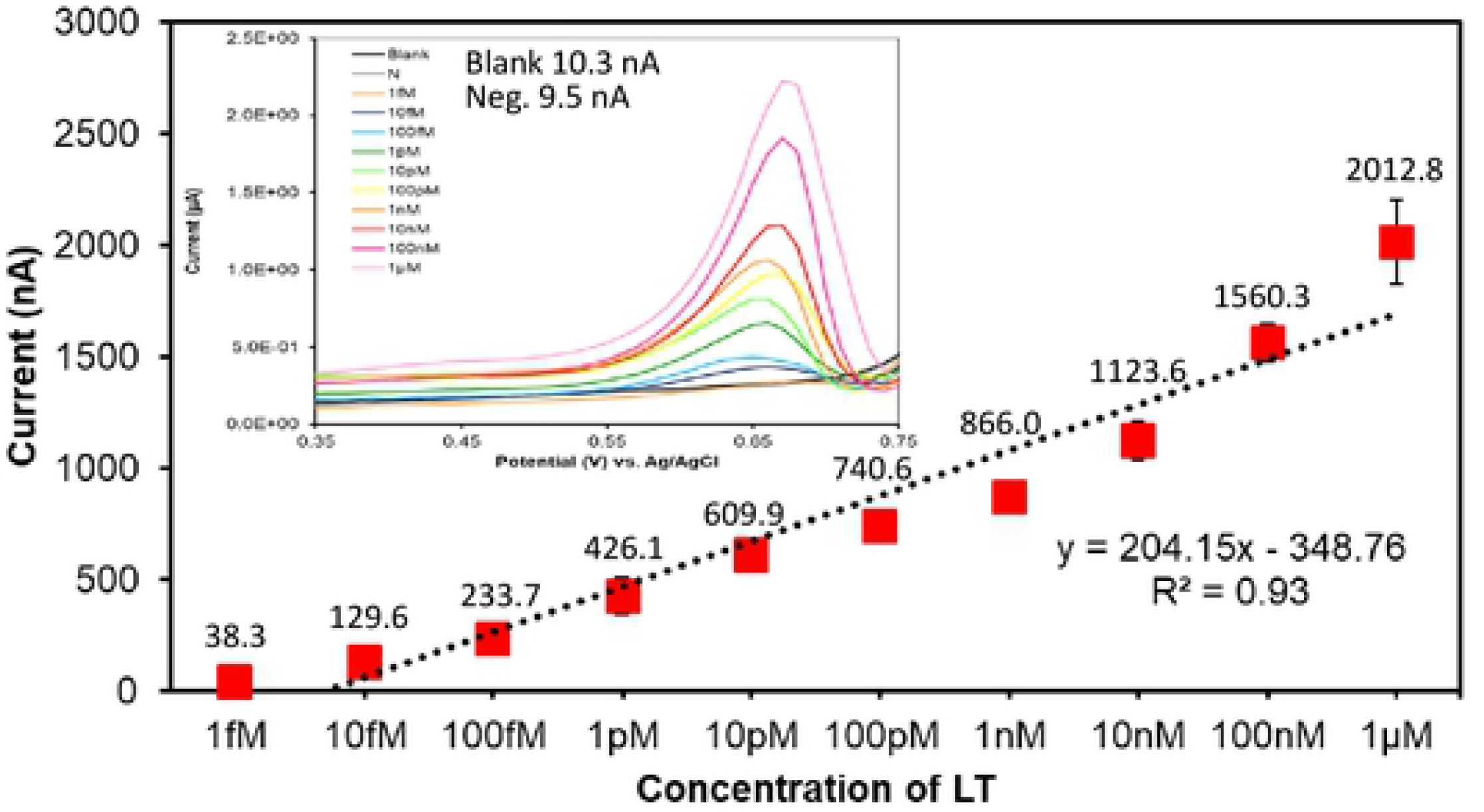
Calibration curve of synthetic wAlbB linear target strand displacement platform using electrochemical detection.

**Figure 5.** Electrochemical detection of wAlbB LAMP products (L) and PCR products (P). PCR reactions were performed for the general *wsp* gene (81F/691R), *wAlbA* gene (328F/691R), and *wAlbB* gene (183F/691R). Different mosquito species were included as follows: wild-type *Ae. aegypti* (AegW), wAlbB trans-infected Thai *Ae*. *aegypti* (AegB), and *Ae. albopictus* (Alb). Samples giving peak currents above 30 nA were considered positive.

**Figure 6.** Electrochemical detection of macerated mosquitoes of laboratory colonies (ML) and field samples (MF). Different mosquito species were included as follows: wild-type *Ae. aegypti* (AegW), wAlbB trans-infected Thai *Ae. aegypti* (AegB), *Ae. albopictus* (Alb), *Cx. gelidus (Cx.gel)* and *Cx. vishnui (Cx.vis)*. Samples giving peak currents above 30 nA were considered positive.

Initially, we wanted to use biosensor in combination with LAMP amplification so as to reduce ambiguity in LAMP visual judgement and increase speed of the detection. We had applied this technology to the LAMP amplified samples (Figure 5–L). The results obtained from electrochemical detection clearly showed a good discrimination between wAlbB-infected mosquitoes (L.Alb and L.AegB) and non-infected mosquitoes (L.AegW). For the positive target, PCR products of *Wolbachia* detection with general wsp primers of the mosquitoes containing wAlbB (P.wsp Alb, P.wsp AegB, P.wAlbB Alb, and P.wAlbB AegB) were detected. The detection signal for negative samples from the mosquitoes without wAlbB (P.wsp AegW) were lower than the threshold value at 30 nA, as shown in Figure 5. In addition, electrochemical detection with the wAlbB targeting capture probe did not respond to the PCR product of wAlbA (P.wAlbA Alb); though the mosquito DNA templates contained wAlbB. This test could effectively separate the detection of wAlbA and wAlbB.

As we discovered high sensitivity of *Wolbachia* detection with the electrochemical DNA sensors, we applied it to test the presence of the wAlbB region directly in the macerated mosquito samples without any *in vitro* amplification. Figure 6 demonstrates positive detection for wAlbB trans-infected Thai *Ae. aegypti* (ML.AegB1-2), *Ae. albopictus* (ML.Alb1 and MF.Alb1-2), and *Cx. vishnui* (MF.Cx.vis1-3), as well as the negative detection for wild-type *Ae. aegypti* (ML.AegW1-2 and MF.AegW1-2) and *Cx. gelidus* (MF.Cx.gel1-2) from both laboratory colonies and field samples. As mentioned earlier, most of the studies reported no *Wolbachia* infection in wild-type *Ae. agypti* and superinfection by wAlbA and wAlbB in *Ae. Albopictus*. We obtained the expected results here. Moreover, previous studies reported the infection of wCon belong to the Supergroup B for the *Cx*. v*ishnui* [31, 33–35], and the infection of *Cx. gelidus* belong to Supergroup A [33–35]. Our biosensor platform only detected the signal in *Cx*. v*ishnui* in Supergroup B. Though the detection was not specific to only wAlbB, a combination of the conventional morphological taxonomy and the molecular detection of mosquito species biomarkers [14, 16] could simply differentiate the mosquitoes in *Aedes* and *Culex* groups.

## Discussion

This study developed a LAMP combined with the electrochemical detection of AuRP from a DNA strand displacement platform for detection of the wAlbB strain of *Wolbachia* bacteria. The methods, i.e., LAMP, LAMP plus BIOSENSOR, and BIOSENSOR alone, could be applied as surveillance and monitoring tools for the *Wolbachia* trans-infected *Ae*. aegypti release programs. Since most studies thus far reported an absence of *Wolbachia* in the wild-type *Ae. aegypti* [11, 31, 33, 35–37], it can be assumed that the *Wolbachia* in *Ae. aegypti* detected in the field surveillance and monitoring study is likely to be from the release programs.

Although, there were few previous studies reporting the detection of natural infection of *Wolbachia* in *Ae. aegypti* [38–45], the *Wolbachia* detection methods in these studies employed only molecular approaches, which are prone to contamination and may be subjected to horizontal gene transfer from the adjacent larvae or parasitic nematodes. Only two studies reported the successful establishment of laboratory colonies of *Wolbachia*-infected *Ae. aegypti* and demonstrated the inherited vertical transmission of *Wolbachia* to F2 [13] and F4 [38] generations. In another independent study [11], however, the cytoplasmic incompatibility and molecular detection on the putatively *Wolbachia*-infected *Ae. aegypti* Las Cruces colony (New Mexico) of the previous work [13] were examined, but *Wolbachia* in this colony could not be found. Therefore, the authors concluded that the evidence of *Wolbachia* in *Ae. aegypti* was not compelling [11]. Regarding the intangible evidence of *Wolbachia* infection in natural *Ae. aegypti*, comprehensive monitoring of the infection status of *Wolbachia* should be continued, especially prior to the release of *Wolbachia* trans-infected mosquitoes. Our detection schemes using LAMP, BIOSENSOR, or a combination could serve this purpose well, as these methods are much more sensitive than the conventional PCR method and can reduce the need for laboratory equipment and molecular biology specialists.

Regarding the cost analysis, excluding DNA extraction, LAMP reagents would cost around $1.5–$3.0/reaction [14, 16]. PCR costs around $0.7. The qPCR cost around $1.0 . If the qPCR included triple replicates and standard curve preparation, the cost could be increased to $3-5 per sample [16]. For BIOSENSOR, the approximate cost would be $2.0. The crude DNA extraction used in this study costed less than $0.5 per sample; although the DNA extraction kit might cost up to $10 per sample. In addition, LAMP required only a single temperature controlled heat block with UV cabinet or clean space, at an investment cost around $1,200. PCR will cost at least $5,500 with a simple gel visualizing instrument, while qPCR will be at least $40,000. Although the electrochemical detection of BIOSENSOR costs around $1,300 in the start-up experiment, the method is cost-effective in the long term. Needless to say, both LAMP and BIOSENSOR, including the combination, would greatly increase the speed of *Wolbachia* detection. Therefore, these methods are very suitable for application in the field, where assessment of expensive molecular laboratory instruments is limited.

We tested the stability of reagent mixture (- template) storaging in the freezer (−20°C) to minimize errors caused by pipetting. Upon adding the DNA template, the LAMP reagent stored up to 20 days could amplify the wAlbB trans-infected Thai *Ae. aegypti*, as indicated by the ladder-like band in agarose gel; although, the faint blue color could be observed when the reagent was stored up to 30 days (Figure S3). It is noteworthy that a number of recent works, and also commercial products, support the possibility of preparing the LAMP reagent in freeze-dried form. Other studies showed that the lyophilized LAMP reagents remained stable for 24 months when stored at 4°C, 28 days at 25°C, 20 days at room temperature, and 2 days at 37°C [39, 40]. In addition, it is also possibile to prepare the strand displacement biosensor reagent in the lyophilized prehybridization mixture. A previous study demonstrated that the prehybridization mixture stored at 4°C is stable up to 3 months without significant decrease in the current signal [24]. However, a decrease of 18% and 30% in the current signal was found in the mixture stored at 25°C and an outdoor ambient temperature (24–34°C) for 50 days, respectively. Further study is needed to apply the lyophilization technique to the *Wolbachia* detecting reagent so as to facilitate the studies which have limited resource settings.

The storage period of the dead mosquito body was also an important concern for a field survey. The results showed that the LAMP reaction could amplify 2 among 3 mosquito samples kept in a freezer (−20°C, without ethanol) up to 10 days (Figure S4). Storage of mosquito samples at 4°C, 27°C, and 37°C for 10 days did not produce the ladder-like bands. However, the LAMP color development results of 2–3 samples of 10-day samples stored at 4°C, 27°C, and 37°C gave blue color. The results from other days were false positive as compared to the negative control (wild-type *Ae. aegypti*). This is consistent with the previous work, which reported that 6 among 10 mosquito samples kept at −20°C for 7 days gave a positive color as compared to the color of the no-template-control [14]. For the samples kept at 14 days and 21 days, 5 and 1 mosquito samples, respectively, among the 10 total samples each were positive. A LAMP positive signal of 2 mosquito samples among 10 samples, which had been stored at 4°C, could be observed up to 2 weeks, while the signal at 37°C could not be observed from 1-week samples [14]. Since the negative (no-*Wolbachia* mosquito control) was not prepared, the results reported might be overestimated. We also observed clear distinguishable colors between no template controls and mosquito samples up to 30 days from samples stored at 37°C. Interestingly, PCR gave positive results for the dead mosquitoes kept at −20°C for at least 30 days and 4°C for 20 days (Figure S5), although the LAMP could not detect any signal. This might be due to degradation of DNA. With respect to mosquito storage conditions, detection of *Wolbachia* from dead mosquitoes stored in a dry condition up to 30 days at 26°C and 10 days at 37°C has been reported [16]. The use of the Genie1 III machine with real-time fluorescence detection was found to increase the sensitivity and reliability of typical LAMP detections with gel electrophoresis and color development [16]. The speed of detection could be increased to 6–12 mins using 6 LAMP primers (including 2 loops) [16].

The LAMP primers and electrochemical biosensing method with strand displacement platform were successfully employed to detect the mosquito samples containing the wAlbB strain of *Wolbachia* bacteria. The tests provided high sensitivity and specificity suitable for field surveys of mosquito distribution in *Wolbachia*-based projects using wAlbB trans-infected *Ae*. *aegypti* and the monitoring of natural *Wolbachia* infections in the wild-type *Ae*. *aegypti*. This knowledge will have tremendous impact, enhance the field of biological control of mosquito vectors, and reduce the problems of arboviral transmission causing millions of deaths in world populations annually.

## Acknowledgements

The authors would like to acknowledge Dr. Lee Su Yin from AIMST University; Dr. Surang Chankhamhaengdecha, Mr. Thanawat Sridapan, Miss Nuanla-ong Kaeothaisong from Mahidol University for their technical assistance; and Mr. David Blyler for English editing. The authors declared that there is no conflict of interest.

## Supporting information captions

**Figure S1.** Detection of *Wolbachia* wAlbB gene using LAMP assays with Ethidium bromide stained gel (A) and Hydroxy Naphthol Blue indicator (B) under different DNA template mass of the different mosquito species including *Aedes albopictus* (1), *Aedes aegypti* (2), and *Wobachia* transinfected Thai *Aedes aegypti* (3). (N) is no-template control and (M) is Invitrogen™ 1 Kb Plus DNA Ladder. The sizes of DNA (bp) were indicated. LAMP reaction was performed using 3.2 units of *Bst* 2.0 DNA polymerase at 65°C for 90 min.

**Table S1.** Detection of *Wolbachia* wAlbB gene using LAMP assays with Hydroxy Naphtol Blue indicator (B) under different *Bst* 2.0 Polymerase concentrations of the different mosquito species. N is no-template control. LAMP reaction was performed at 65°C for 90 min.

**Figure S2.** Detection of *Wolbachia* wAlbB gene using LAMP assays with Ethidium bromide stained gel (A) and Hydroxy Naphthol Blue indicator (B) under different incubation times for 60 and 90 min with a DNA template of 20 and 40 ng. P is *Aedes albopictus* and N is no-template control. M is Invitrogen™ 1 Kb Plus DNA Ladder. The sizes of DNA (bp) were indicated. LAMP reaction was performed using 3.2 units of Bst 2.0 DNA polymerase at 65°C for 90 min.

**Figure S3.** Detection of *Wolbachia* wAlbB gene using LAMP assays with Ethidium bromide stained gel (A) and Hydroxy Naphthol Blue indicator (B) with varying LAMP reagent (included *Bst* polymerase) at −20°C freezer under different incubation times for 60 and 90 min with a DNA template of 20 and 40 ng. P is *Aedes albopictus* and N is wild-type *Ae. agypti*, and NT is no-template control (NTC). M is Invitrogen™ 1 Kb Plus DNA Ladder. The sizes of DNA (bp) were indicated. LAMP reaction was performed using 6.4 units of Bst 2.0 DNA polymerase at 65°C for 60 min and 80°C for 10 min.

**Figure S4.** Detection of the *Wolbachia* wAlbB gene using LAMP assays with Ethidium bromide stained gel (upper) and Hydroxy Naphthol Blue indicator (lower), with the dead mosquitoes stored at different temperatures including −20°C (A), 4°C (B), 27°C (C), and 37°C (D). 1–3 are wAlbB infected *Ae. agypti* mosquitoes. P is *Aedes albopictus*. N is wild-type *Ae. agypti*, and NT is no-template control (NTC). M is Invitrogen™ 100 bp or 1kb plus DNA Ladder. The sizes of DNA (bp) were indicated. LAMP reaction was performed using 6.4 units of *Bst* 2.0 DNA polymerase, 65°C for 60 min and 80°C for 10 min.

**Figure S5.** Detection of *Wolbachia* wAlbB gene using PCR with Ethidium bromide stained agarose gel, with the dead mosquitoes stored at different temperatures including −20°C, 4°C, 27°C, and 37°C. 1–3 are wAlbB infected *Ae. agypti* mosquitoes. P is *Aedes albopictus*. N is wild-type *Ae. agypti*, and NT is no-template control (NTC). “-” is no loading well. M is Invitrogen™ 100 bp DNA Ladder. The sizes of DNA (bp) were indicated.

